# Contrast-enhanced sonography with biomimetic lung surfactant nanodrops

**DOI:** 10.1101/2020.11.03.367235

**Authors:** Alec N. Thomas, Kang-Ho Song, Awaneesh Upadhyay, Virginie Papadopoulou, David Ramirez, Richard K. P. Benninger, Matthew Lowerison, Pengfei Song, Todd W. Murray, Mark A. Borden

## Abstract

Nanodrops comprising a perfluorocarbon liquid core can be acoustically vaporized into echogenic microbubbles for ultrasound imaging. Packaging the microbubble in its condensed liquid state provides distinct advantages, including *in situ* activation of the acoustic signal, longer circulation persistence, and the advent of expanded diagnostic and therapeutic applications in pathologies which exhibit compromised vasculature. One obstacle to clinical translation is the inability of the limited surfactant present on the nanodrop to encapsulate the greatly expanded microbubble interface, resulting in ephemeral microbubbles with limited utility. In this study, we examine a biomimetic approach to stabilizing an expanding gas surface by employing the lung surfactant replacement, Beractant. Lung surfactant contains a suite of lipids and surfactant proteins that provides efficient shuttling of material from bilayer folds to the monolayer surface. We therefore hypothesized that Beractant would improve stability of acoustically vaporized microbubbles. To test this hypothesis, we characterized Beractant surface dilation mechanics and revealed a novel biophysical phenomenon of rapid interfacial melting, spreading and re-solidification. We then harnessed this unique spreading capability to increase the stability and echogenicity of microbubbles produced after acoustic droplet vaporization for *in vivo* ultrasound imaging. Such biomimetic lung surfactant-stabilized nanodrops may be useful for applications in ultrasound imaging and therapy.

**Graphical Abstract:** 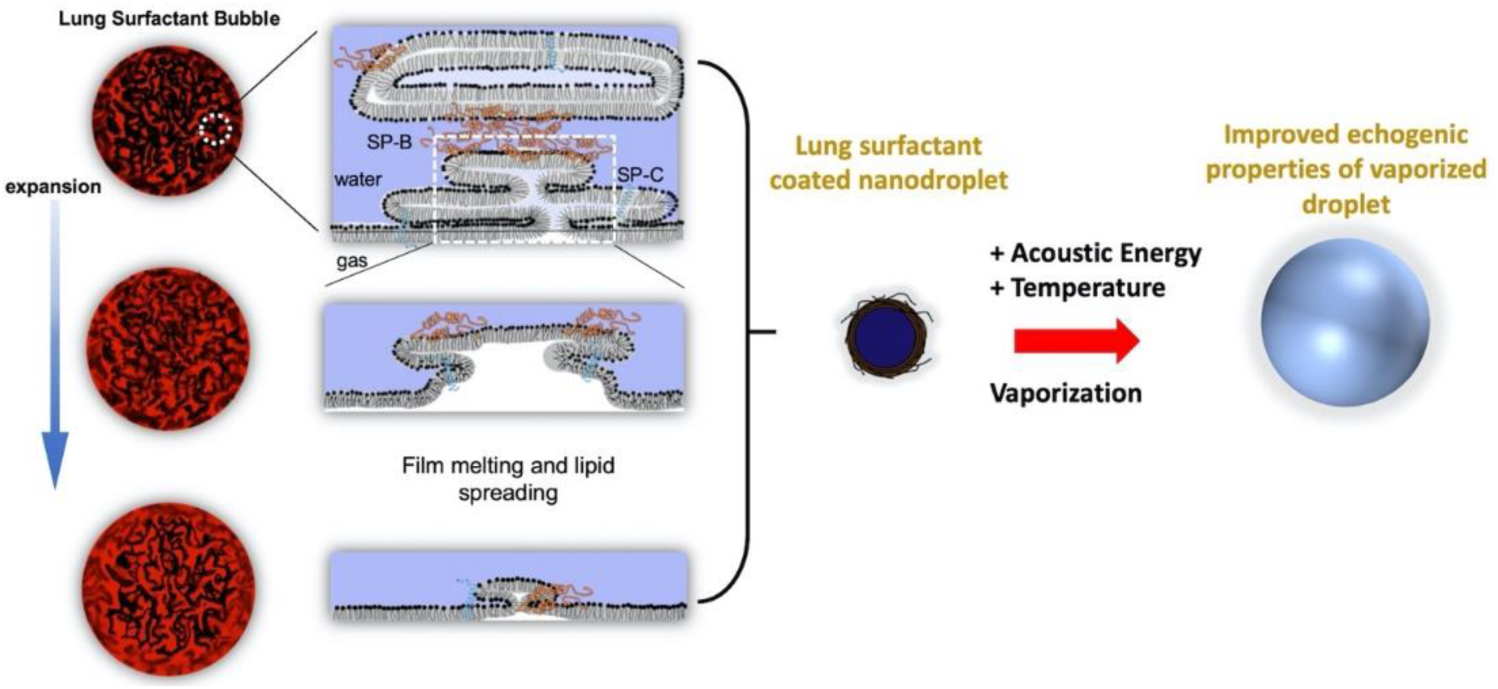

## 1. Introduction

Ultrasound is the most widespread clinical imaging modality today. The advent of smartphone-controlled imaging probes, super-resolution imaging techniques, and commercial interest in targeted non-invasive ultrasonic therapies continues to signal ultrasound’s growing relevance in advanced imaging and image-guided therapy [1]. Microbubbles, commonly 1-10 µm diameter lipid-coated gas spheres, are particularly echogenic structures well-suited for use as ultrasound contrast agents. Due to their size, however, microbubbles may be rapidly trapped by size-dependent filtering organs, such as the lung, liver and spleen [2], and are unable to transport outside of vascular compartments, limiting contrast-enhanced ultrasound (CEUS) assessment to specific luminal and endothelial features. In imaging applications for cancer [3], heart disease [4] and diabetes [5], the promise of extravascular CEUS offers imaging of significant pathophysiological indicators, namely the formation and prevalence of inflammation and neovasculature, or vascular leakage into interstitial tissue, which may enable diagnosis and indicate treatment efficacy. Vaporizable nanodrops also provide excellent activatable or “blinking” probes for super-resolution ultrasound [6], which can generate exquisite images of the microvasculature deeper in tissue than can be achieved with current optical microscopy techniques [7].

During sonography, nanodrops form echogenic microbubbles via acoustic droplet vaporization (ADV) [8], a process in which a brief increase in acoustic amplitude triggers rarefaction-induced homogeneous nucleation of a critical vapor embryo—the beginning of bubble formation [9]. ADV is triggered using clinical medical ultrasound imaging systems and can be conducted in deep tissue. Recent work has focused on engineering the fluorocarbon core of nanodrops for ADV at clinically relevant ultrasonic mechanical index (MI) [10,11].

Implementation of acoustically activated nanodrops faces the challenge of microbubble instability due to insufficient shell coverage. Droplet vaporization typically produces a 25-fold surface area expansion [12], which contributes to the failure of conventional surfactant shells to fully encapsulate and stabilize the resultant microbubble against coalescence and rapid dissolution [13]. The transience of the nanodrop-turned-microbubble significantly reduces the utility of such nanodrops in imaging applications beyond proof-of-concept.

Here, we investigate a biomimetic approach employing the lung surfactant replacement, Beractant (Survanta, AbbVie, North Chicago, IL), to overcome the problem of limited film coverage on the vaporized microbubble. Much like the encapsulating lipid of microbubble shells, the role of surfactant in the lungs is to reduce the surface tension of lung alveoli—the primary microstructural mediators for gas exchange in the lungs. Beractant has been shown to stabilize microbubbles by forming a monolayer at the gas/water interface with attached bilayer folds that extend into the aqueous phase [14]. The Beractant film is transported from the monolayer to the bilayer folds during hydrostatic compression of the microbubble [15]. These attached bilayer folds may serve as a film reservoir for the expanding interface. We therefore hypothesized that Beractant may readily deploy with the vaporizing nanodrop to stabilize the resulting microbubble.

Prior to examining ultrasound imaging utility of Beractant-coated nanodrops *in vivo*, we investigated the film dilatational properties *in vitro* using a laser-acoustic method and discovered a unique interfacial melting and spreading phenomenon by Beractant [16]. We then examined the echogenicity and ultrasound contrast longevity *in vitro* and *in vivo*. The results support our hypothesis that biomimetic lung surfactant provides improved spreadability to stabilize the vaporized microbubble. Finally, we explored the potential for super-resolution imaging in an *ex ovo* chorioallantoic membrane (CAM) and for imaging inflamed endocrine pancreas vasculature *in vivo* in a mouse model of type-1 diabetes.

## 2. Methods and Materials

### 2.1. Microbubble Synthesis and Characterization

#### 2.1.1. Lung Surfactant (Beractant) Microbubbles

Lung surfactant microbubbles were made using the intratracheal lung surfactant replacement therapy Beractant (Survanta®, Abbvie Inc, North Chicago, IL). Beractant is a highly concentrated aqueous mixture of lung surfactant proteins and lipids extracted from excised bovine lungs and supplemented with synthetic DPPC, palmitic acid and tripalmitin. The biochemical composition of Beractant is reported in the package insert as 25 mg/mL phospholipids (mainly DPPC), 0.5-1.75 mg/mL triglycerides, 1.4-3.5 mg/mL free fatty acids and less than 1.0 mg/mL protein (SP-B, SP-C). The surfactant composition is somewhat altered from the native surfactant by the extraction process, with reduced surfactant proteins SP-B and SP-C, and elimination of cholesterol, SP-A and SP-D. Beractant was stored at 4 °C, and care was taken to maintain and monitor sterility. Microbubbles coated with Beractant were prepared as previously described [14]. Briefly, 1.92 mL PBS was added to 80 µL of Beractant, for a total concentration of 1 mg/mL, in a 3-mL glass serum vial. The vial was capped and sealed, and bath sonicated at 40 °C for 20 min to disperse the suspension. The vial was cooled to room temperature, and the air headspace was exchanged with perfluorobutane (PFB) gas. The vial containing Beractant suspension and PFB gas was mechanically activated using a dental amalgamator (VialMix) for 45 s resulting in a milky white suspension containing microbubbles. The vial of microbubbles was then cooled in an ice bath. The suspension was then extracted with a 3-mL syringe, and microbubbles were isolated by 4 centrifugation wash cycles at 40 relative centrifugation force for 1 min (Eppendorf 5804, Hamburg, Germany), where the infranatant was discarded and the microbubble concentrate was saved and resuspended.

#### 2.1.2. DPPC Microbubbles

Control microbubbles were made with the pure lipid component 1,2-dipalmitoyl-sn-glycero-3-phosphocholine (DPPC) (Avanti Polar Lipids, Alabaster, AL), as previously described [15]. DPPC was weighed and dissolved in chloroform (Sigma-Aldrich, St. Louis, MO). The lipid solution was then vacuum dried for 4 hours to remove the organic solvent, leaving behind a transparent film. The dried transparent film was rehydrated in 0.2-µm filtered, 10 mM phosphate buffer saline (PBS) to a total lipid concentration of 4.0 mg/mL. The lipid suspension was then heated at 50 °C, stirred, and sonicated at 3/10 power using a half-inch probe tip submerged below the water surface for 3 min (Branson 450 Sonifier, Danbury, CT). The suspension was cooled to ambient temperature and 2.0-mL aliquots were distributed to 3-mL glass serum vials. The vials were capped and sealed, and the air headspace was replaced with perfluorobutane gas (Fluoromed, Round Rock, TX). The vial containing lipid suspension and perfluorobutane (PFB) gas was mechanically activated using a dental amalgamator (VialMix, Bristol-Myers-Squibb, NY) for 45 s resulting in a milky white suspension containing microbubbles. The vial of microbubbles was then cooled in an ice bath.

#### 2.1.3. Perflutren Lipid Microbubbles

Microbubbles were made using the same shell composition as for the commercially available microbubble Perflutren lipid microspheres (Definity®, Lantheus Medical Imaging, N. Billerica, MA), which is approved for contrast-enhanced ultrasound imaging by the U.S. Food and Drug Administration. Briefly, a lipid solution containing 1,2-dipalmitoyl-sn-glycero-3-phosphocholine (DPPC) (Avanti Polar Lipids, Alabaster, AL), 1,2-diplamitoyl-sn-glycero-3-phosphate (DPPA) and 1,2-dipalmitoyl-sn-glycero-3-phosphoethanolamine-N-[azido(polyethylene glycol)-5000 (DPPE-PEG5000) (NOF America, White Plains, NY) were used in a molar ratio of 82:10:8, respectively. Lipids were weighed and dissolved in chloroform (Sigma-Aldrich, St. Louis, MO). The lipid solution was then vacuum dried for 4 hours to remove the organic solvent, leaving behind a transparent film. The dried transparent film was rehydrated in 0.2-µm filtered, 10 mM phosphate buffer saline (PBS) to a total lipid concentration of 0.75 mg/mL. The lipid suspension was then heated at 50 °C, stirred, and sonicated at 3/10 power using a half-inch probe tip submerged below the water surface for 3 min (Branson 450 Sonifier, Danbury, CT). The suspension was cooled to ambient temperature and 2.0-mL aliquots were distributed to 3-mL glass serum vial. The vials were capped and sealed, and the air headspace was replaced with perfluorobutane gas (Fluoromed, Round Rock, TX). The vial containing lipid suspension and perfluorobutane (PFB) gas was mechanically activated using a dental amalgamator (VialMix, Bristol-Myers-Squibb, NY) for 45 s resulting in a milky white suspension containing microbubbles. The vial of microbubbles was then cooled in an ice bath. The filling gas of PFB was used instead of octofluoropropane, which is used in the commercial product, Perflutren lipid microspheres. This different filling gas was required to produce a more stable nanodrop emulsion.

#### 2.1.4. Nanodrop Synthesis

Nanodrops were prepared as described by Mountford et al. [11,17,18]. The vial containing either Perflutren lipid or lung surfactant (Beractant) microbubbles was cooled in an isopropanol bath to −10 °C under continuous mixing to avoid freezing. After ∼1 min, the microbubble suspension was pressurized to 170 kPa gauge pressure by inflowing PFB gas into the vial. The pressurization by PFB gas resulted in conversion of the milky white microbubbles suspension to a transparent nanodrops suspension.

#### 2.1.5. Microbubble and Nanodrop Sizing and Counting

Microbubbles were sized with a Coulter Multisizer III (Beckman-Coulter, Brea, CA) using a 30-μm aperture set to measure particles between 0.8 and 18 μm in diameter. Immediately after synthesis, microbubbles rested for 10 min before the suspension was mixed and diluted to 0.2% v/v in Isoton (Beckman-Coulter). For each vial, three measurements were taken and background measurements of Isoton were subtracted from the microbubble measurements (supplementary Fig. S3). Nanodrop size distribution and concentration were not directly measured. As a first approximation, the nanodrop concentration was assumed to be identical to the microbubble concentration assuming that there was no loss during the nanodrop synthesis process. Additionally, post-ADV microbubble size and concentration was not measured owing to instability from handling and processing, the cardiovascular circuit, dynamics of sonication, and the open thermodynamic system that allowed gas transfer between the atmosphere.

### 2.2. Laser-Acoustic Measurement of Microbubble Resonance Curves

The photoacoustic method described by Lum et al. [16] was used to measure the viscoelastic properties of individual microbubble shells. The sample was held in a temperature-controlled microscope chamber. Microbubbles coated with either lung surfactant or DPPC were pipetted into the 300-µL chamber, and single microbubbles were selected and sized by a dark-field microscopy image. The selected microbubble was then driven into oscillation by photoacoustic pressure waves produced with an intensity-modulated laser source at a wavelength of 1550 nm, where water is strongly absorbing at that wavelength. The driving infrared laser was positioned near (∼100 *μm*) the bubble (Fig. 1a), and the frequency of the laser intensity modulation was swept from 0.5 to 3.0 MHz with a function generator in 50 kHz steps. A second tracking laser operating at 488 nm was focused onto the microbubble, and the transmitted illumination was detected with a photodetector. The output of the photodetector was sent to a lock-in amplifier to detect the magnitude of the response at the modulation frequency and allowing for the acquisition of a resonance curve for the target microbubble. The position and width of the resonance curve was used to determine the shell elasticity and viscosity, respectively, using the linearized Chatterjee-Sarkar model [19] (see Supplemental section describing theory and calculations).We first examined the shell elasticity and viscosity as a function of temperature (22 to 47 °C) on microbubbles that had equilibrated for at least 40 min at the desired temperature. For each temperature point, n ≥ 20 microbubbles were tested. We next studied the transient effects of microbubble growth and dissolution on the shell elasticity and viscosity. For these experiments, the initial state of the microbubble was measured at room temperature (∼25 °C) with continuous measurements for ∼20 min. The sample was then heated at a rate of 0.9 ± 0.08 °C/min by two resistive heaters adhered to the chamber until reaching a constant temperature of ∼37 °C. Measurements were continuously recorded for up to an hour after reaching 37 °C during which time the microbubbles were observed to grow and then shrink back to their initial size. Shell viscosity and elasticity were measured on individual microbubbles (n=16 for Beractant and n=13 for DPPC) during the expansion and compression excursions.

**Fig. 1.**
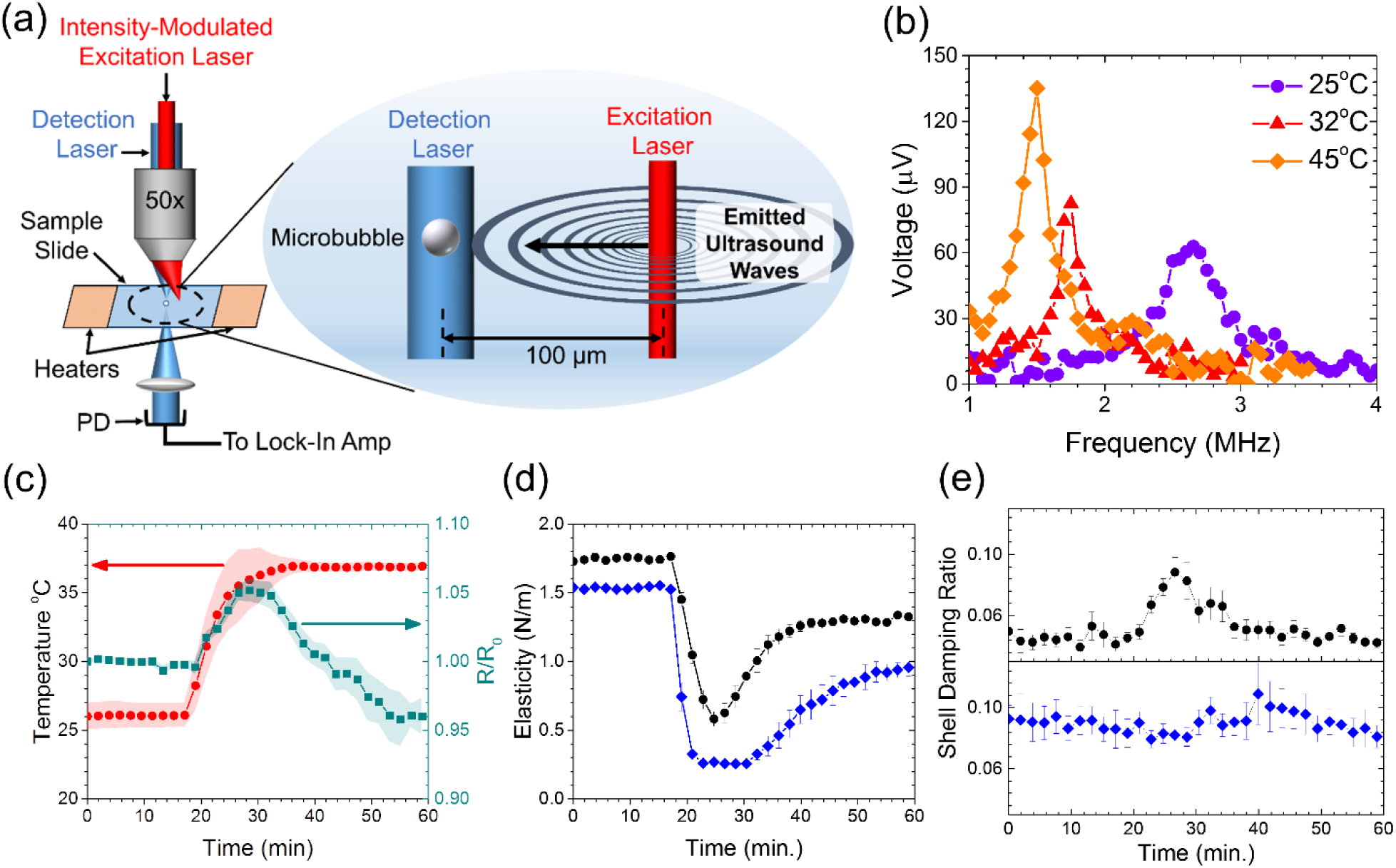
Viscoelastic properties of the microbubble shells. (a) Illustration of the laser-acoustic technique used to drive microbubble oscillations by photoacoustic sound waves emitted from absorption of nearby excitation laser pulses. The oscillations were detected by the photodetector (PD) through forward light scattering of the continuous wave detection laser. (b) The output voltage from the lock-in amplifier at three different temperatures, representing the magnitude of radial oscillations for Beractant-coated microbubbles taken at different temperatures. (c) The temperature profile (red circles) and normalized radius (green squares) of Beractant microbubbles (n=16). Shaded regions show s.e.m. The average shell elasticity (d) and damping ratio (e), a measure of surface viscosity, for Beractant (black circles, n=16) and DPPC (blue diamonds, n=13) films during the bubble growth and restabilization process. The simultaneous rise in viscosity and drop in elasticity for Beractant is a hallmark for interfacial melting and spreading.

### 2.3. In vitro Acoustic Nanodrop Vaporization Experiment

A volume of 5-mL of diluted lung surfactant nanodrops or Perflutren lipid nanodrops (5 × 10^7^per mL) were placed into a section of cellulose tubing (Spectrum Chemical, New Brunswick, NJ) that was submerged in a water bath maintained at 37 °C. The tubing contained a small 2-mm magnetic stir bar to keep the tube well-mixed. The tube was positioned at the center of the field of view of the Acuson 15L8 ultrasound probe attached to an Acuson Sequoia C512 clinical ultrasound imaging scanner (Siemens, Munich, Germany). The nanodrops were allowed to equilibrate for 5 min. The focal depth was set to 20 mm, the mechanical index was turned down to its lowest value, and the center frequency was fixed to 8 MHz. All ultrasound image sequences were acquired with a constant dynamic range for all experiments (75 dB). After thermal equilibration, the video images from the scanner were recorded by a LabVIEW program (National Instruments, Austin, TX) set to acquire an image every 250 ms. The MI was then rapidly increased to the desired peak MI. For example, initially the MI was set to 0.04 and then quickly increased to 0.63 MI for the remainder of the experiment. Then the cellulose tubing was flushed and new nanodrops entered the focal region before commencing a subsequent experiment. Based on preliminary experiments, no vaporization for either shell type was observed to occur below an Output Display Standard (ODS) MI of 0.30 defined on the Acuson Sequoia. Images were recorded for 3 min after the peak MI was reached. Ultrasound images were digitized using a custom MATLAB (Mathworks) script to calculate the video intensity in the region of interest of each time-stamped image. Accordingly, the ODS MI was corrected by the following equation:

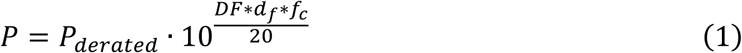

where *P* is the peak negative pressure (PNP) in water, *p*_*derated*_ is the PNP derated for tissue attenuation (as read by the ODS), *DF* is the derating factor 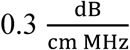 for tissue, *d*_*f*_ is the focus depth of the transducer and *f*_*c*_ is the operating fundamental frequency of the transducer. The actual MI encountered by the nanodrops is a factor of ∼1.7 higher than the ODS MI.

### 2.4. In vivo *Acoustic Nanodrop Vaporization Experiment in the Mouse Kidney*

#### 2.4.1. Animal Preparation

The *in vivo* studies were conducted in accordance with the protocols used by Institutional Animal Care and Use committee at the University of Colorado Anschutz medical campus. All mice were purchased from Jackson Laboratories (Bar Harbor, ME). 10-week-old female C57BL/6J mice were housed in AAALAC approved animal care facilities in compliance with PHS policy. The mice had *ad libitum* access to drinking water and standard pelleted chow. The feed was no less than 23% protein, 4.5% fat, and 6% fiber. At the time of the scan, the mice weights ranged from 22 to 28 g. Animals were divided into two experimental groups: Beractant nanodrop bolus injection (n=4) and Perflutren lipid nanodrop bolus injection (n=4). Mice were anesthetized via inhaled isoflurane (2%) mixed with oxygen as the carrier gas. Hair around the abdomen was removed using depilatory cream, and the area was cleaned thoroughly using alcohol swab. Mice were kept on a heating pad with their legs secure using surgical tape so as to avoid any movement during the study. Coupling gel was placed on the abdomen area for the ultrasound imaging probe. A custom 27-gauge winged infusion set was inserted in the tail vein and secured with VetBond tissue adhesive (3M Animal Care Products, St. Paul, MN).

#### 2.4.2. Image Acquisition

The same Siemens Acuson Sequoia C512 scanner and 15L8 probe was used to image the kidney, chosen as the organ of interest due to its high blood perfusion. Once the longitudinal view of the kidney was located using high frequency B-mode (14 MHz), the transducer was secured in place by a clamp before changing the imaging settings to acquire in contrast-enhanced as follows. Contrast acquisitions were imaged in dual B-mode and contrast mode, with the focal depth set to corresponding to the depth of the kidney, at a frequency of 8 MHz and a fixed MI of 0.7, shown from *in vitro* studies to result in maximum enhancement; Cadence Pulse Sequencing (CPS) gain was set to 0 dB and the dynamic range was set at 50 dB. A high MI flash pulse was not necessary to induce vaporization. The mice were then injected with a bolus dose of 200 µL of nanodrops at a concentration of 1 × 10^9^ per mL. IC Capture 2.4 software (The Imaging Source LLC, Charlotte, NC) was used to acquire image video data from the ultrasound scanner, starting right before the nanodrop injection and for 5 min post-injection, which was longer than necessary to ensure that the contrast enhancement returned to baseline levels prior to injection. The videos were then analysed for average contrast intensity within a region of interest (ROI) in the kidney to generate time-intensity curves. These curves were then analysed with a first-order elimination pharmacokinetic model to determine the half-life and area-under-the-curve (AUC) using a custom MATLAB code.

#### 2.4.3. Image and Data Analysis

The acquired videos were post-processed off-line using custom MATLAB code to determine contrast enhancement generated by the nanodrops. An ROI was drawn on the first frame of each video around the kidney in the B-mode and transposed to the CPS part of the dual B-mode and CPS image. The average pixel intensity within this ROI on CPS was then calculated for all frames. Baseline intensity (pre contrast injection) was taken as the average of the ROI mean over the first 15 frames (0.43s) of the video. Baseline-subtracted, mean ROI intensities were plotted against time and used to calculate contrast enhancement and persistence. Statistical comparisons between groups were performed in GraphPad Prism 8.2 (GraphPad Software, La Jolla, CA, USA). Statistical significance was set *a priori* at p < 0.05 and all data is presented as mean ± standard deviation unless otherwise stated. First, the AUC was calculated and compared between the two groups (Beractant and Perflutren lipid) via a t-test using the mean AUC, standard of the error mean (SE) and the degrees of freedom of the AUC fitting for each group. Then, for the Beractant nanodrops resulting in considerable contrast enhancement, the half-time for contrast persistence was calculated by fitting a mono-exponential decay model to the curve from its highest intensity point. Since contrast enhancement was short-lived in all cases and to negate subsequent motion artefacts, the time-series used in analysis was capped at 90 s post injection. All contrast enhancement was confirmed to be back to baseline levels prior to that point. Finally, as a demonstration of a potential imaging application, a maximum intensity projection (MIP) of all contrast enhanced frames was performed after localization of microbubble positions. For each frame after nanodrop injection, the intensities of all pixels were baseline-corrected by subtracting the intensities of the corresponding pixels from the first frame prior to nanodrop injection, before binarizing the resulting image by histogram thresholding. Finally, all frames were summed, resulting in a MIP of the kidney ROI over time after nanodrop injection.

### 2.5. Ultrasound Imaging of Nanodrop Distribution in Chicken Embryos

#### 2.5.1. Embryo preparation

Fertilized chicken eggs were obtained from the University of Illinois Poultry Research Farm and placed into a humidified rocking incubator (Digital Sportsman Cabinet Incubator 1502, GQF) at a temperature between 37-38 °C, with passive humification via a water pan that was refilled daily. Eggs were turned automatically every 2 hours. On the fourth day of embryonic development *ex ovo* chorioallantoic membrane (CAM) assays were generated by opening the eggshell with a rotary Dremel tool and transferring egg contents into plastic weigh boats. *Ex ovo* CAMs were housed in a separate humidified incubator (Darwin Chambers HH09-DA) held at 37-38C with a relative humidity maintained at 90% until ready for ultrasound imaging.

#### 2.5.2. Image Acquisition

*Ex ovo* CAMs were removed from their housing incubator and placed onto a heating pad. Prior to ultrasound imaging a glass capillary needle was attached to 8 cm of Tygon R-3603 laboratory tubing and fitted onto an 18G 1½” beveled needle with a 1-mL syringe. The glass needle was placed into a CAM surface vessel with the aid of a dissecting microscope (Nikon SMZ800N), and a 70-uL bolus of nanodrops was injected into each chicken embryo. A Verasonics Vantage 256 (Verasonics Inc., Kirkland, WA) ultrasound system equipped with an L35-16vX high-frequency linear array transducer (Verasonics Inc., Kirkland, WA) was used to image the chicken embryos. Ultrasound contact gel was applied to the surface of the transducer to serve as an acoustic stand-off. Plane wave imaging with a pulse-repetition-frequency (PRF) of approximately 33,333 Hz was performed with 9-angle coherent compounding [20] (−4 degrees to 4 degrees, 1-degree step angle), a center frequency of 20 MHz, and an effective PRF (i.e., post-compounding PRF) of approximately 3700 Hz. During the experiment, no-operation time delay was added to slow down the effective frame rate to 50 Hz to allow capturing of nanodrop distribution and movement. The elevational plane thickness of the L35-16vX transducer is approximately 0.8 mm. The ultrasound transducer was coupled either to the head of the chicken embryo to image the brain, or to the side of the weigh boat to produce images of the CAM surface vessels. A custom acquisition script was programmed to step-up the transmit voltage of the acoustic waveform during imaging to acquire both pre- and post-vaporization images. In-phase quadrature (IQ) datasets were collected containing 1500 ultrasound frames (500 for each transmit voltage), corresponding to a 30-s data acquisition length. The 3 voltages used in the Verasonics acquisition were 6 volts (low), 25 volts (medium), and 40 volts (high). The 500 middle voltage frames were not used in the analysis – only as part of an initial voltage ramp-up for vaporization events. For the last 500 frames (high voltage), a single value decomposition (SVD) filter was applied to extract stationary vaporized nanodrops by exploiting their high spatiotemporal coherence. The SVD-filtered data was then accumulated along the temporal dimension. For SVD filtering, the applied values corresponded to removing all velocities below approximately 0.246 mm/s for flow imaging [21]. The SVD filtering used to retain stationary vaporized nanodrops excluded data above this velocity, as well as excluded the lowest singular value (around the 0.005 mm/s to 0.006 mm/s range).

#### 2.5.3. Image and Data Analysis

Ultrasound IQ was divided into two subsets: the first 500 frames (lowest voltage) were used to generate vascular images, and the last 500 frames (highest voltage) were used to localize vaporized nanodrops. In both cases the 3D IQ dataset was reshaped into a 2D Casorati matrix prior to SVD decomposition. A spatiotemporal SVD-based clutter filter was applied to the vascular IQ dataset to reject background tissue signal [22]. This cut-off threshold was determined by the decay rate of the singular value curve [21] and was set to filter out the first 12 singular values in this dataset. Ultrasound microvessel perfusion images were then generated by accumulating the resulting vascular signal along the temporal direction. The IQ dataset containing ADV nanodrops (the last 500 IQ frames at the highest transmit voltage) was also SVD filtered, in this case to extract the stationary nanodrops that had a high spatiotemporal coherence. Two thresholds were empirically determined and applied: a low-order threshold was used to reject tissue, and a high-order threshold was applied to reject all signal components with movement (blood flow, nano/microbubble flow). The lower threshold was set to exclude the first two singular values, and the higher threshold excluded everything above ten singular values.

### 2.4 Ultrasound Imaging in a Diabetic Animal Model

#### 2.6.1. Animal Preparation

All experiments were performed with ethical approval and within guidelines established by the Institutional Animal Care and Use Committee of the University of Colorado Anschutz medical campus. 10-week-old female NOD/ShiLtJ (non-obese diabetic, NOD) mice were purchased from Jackson Laboratories (Bar Harbor, ME). Blood glucose concentrations were monitored weekly for each mouse with a glucometer (Bayer).

#### 2.6.2. Image Acquisition

All mice were anesthetized with isoflurane inhalation for the duration of the experiment (approximately 35 min). Prior to ultrasound imaging, a 27G ½” winged infusion set (Terumo BCT, Lakewood, CO) attached to polyethylene tubing (0.61 OD x 0.28 ID; PE-10, Warner Instruments) was placed into the lateral tail vein. The abdominal fur was removed using a depilatory cream and then cleaned with alcohol swabs. Ultrasound coupling gel was placed on the abdomen and then the transducer was placed. Electrode foot pads were used to monitor electrocardiogram and respiratory rate, while a transrectal temperature probe read the mice’s body temperature. A VEVO 2100 small animal ultrasound machine (Visual Sonics Fujifilm, Toronto, Canada) was used to capture the ultrasound time courses. A MS250 linear array transducer was used at a frequency of 18 MHz. B-mode imaging was performed prior to contrast imaging to identify the pancreas based on a striated appearance and proximity to the kidney, spleen, and stomach. Following positive identification of the pancreas, contrast mode was initiated and a region of interest was placed around the various organs. Prior to contrast infusion, three minutes of background images were saved (‘pre-infusion’); these were used to normalize the time course so comparisons can be made between all animals. Following infusion, a 25-min pause in imaging to allow nanodrop biodistribution, followed by initiation of image acquisition and acoustic ADV. ADV was initiated by a high mechanical index flash of 1.5. The contrast mode settings were as follows: transmit power 10%, frequency 18 MHz, standard beam width, focus depth 3 mm from abdominal wall, contrast gain of 30 dB, and 2D gain 18 dB, with an acquisition rate of 27 frames per second. There were 500 frames collected per ∼18 sec time course. Each time course was collected at 30 s intervals for 30 min. Six background time courses (3 min) were collected prior to infusion.

#### 2.6.3. Image and Data Analysis

To account for breathing-induced motion artifacts each time course was ‘gated’ using a custom Matlab script. A frame from the time-course where the animal is stable, between breaths, was first selected. The script creates a binary mask from the selected region of interest in the image. These regions of interest were selected to comprise the B-mode and contrast mode images from the experimental time course. The images were then cropped so only the B-mode and contrast mode ROIs are evaluated. The intensity values of the two masks were measured relative to one another and any value that is more than three standard deviations away from the median were excluded. Additional frames were removed manually as determined by visual inspection of the B-mode images for motion. The remaining frames were then retained for subsequent analysis. After gating, regions of interest were placed over the pancreas and kidney organs. VEVO CQ software was used to obtain the measured contrast change in the organs throughout all time courses. The measured contrast intensities were binned to 1-min intervals. All time points were background subtracted and normalized to background (signal – background / background) and plotted to compare fold-changes in contrast signal.

## 3. Results and Discussion

Microbubble stabilization during expansion and recompression is facilitated by melting and spreading of the shell, which can be interpreted from two surface viscoelastic parameters: elasticity and shell damping (viscosity). These parameters were examined in microbubbles as temperature was increased between 25°C and body temperature (37°C) over a 10 min period (Fig. 1b,c). Over this time, thermal expansion and outgassing from the surrounding aqueous medium [16] resulted in an average radial expansion of 5.0 ± 0.1%, corresponding to an average ∼10% increase in surface area. The bubble then shrank to slightly less than its original size, on average, owing to Laplace-pressure driven dissolution as dissolved gases in the microscope chamber equilibrated with the atmosphere. The smaller bubble radius tends to increase the resonant frequency, however at elevated temperature the shell stiffness decreases [16], thus the resonant frequency was nearly unchanged. Measurements of elasticity and damping were taken every 2 min during this process. The viscoelasticity of Beractant was compared to dipalmitoyl phosphatidylcholine (DPPC), the main lipid component of mammalian lung surfactant and Perflutren, a clinical microbubble ultrasound contrast agent. The elasticities of both Beractant and DPPC films were observed to drop at the onset of surface dilation (Fig. 1d). Of note however, was that the elasticity of Beractant rebounded, even as the surface area continued to grow. This rebound was not observed for DPPC until after the microbubble had shrunk back to its initial size. Additionally, the minimum elasticity for Beractant (0.57 ± 0.22 N/m) was two-fold greater than for DPPC (0.23 ± 0.05 N/m).

Next, we observed that the damping ratio—a direct measure of the surface viscosity—did not change significantly in DPPC shells but increased twofold on average in Beractant films over the course of the temperature increase (Fig. 1e). The effect can be observed most prominently on an individual microbubble (Fig. 2a-c). The concurrent rise in surface viscosity and drop in elasticity with dilation strongly suggested that the Beractant film melted, which facilitated spreading of the film over the newly formed interface during expansion (shown schematically in Fig. 2d,e). The effect can also be visualized in the mechanical testing curves, shown here as elasticity *vs*. area, where significant hysteresis indicates plastic deformation for Beractant, while lack of hysteresis denotes brittle fracture for DPPC (Fig. 2f,g). To our knowledge, this melting-to-spreading transition has not been previously observed for lung surfactant films. The Beractant film re-solidified upon compression, as indicated by the rise in elasticity and drop in viscosity. It was also observed that *static* Beractant films have higher elasticity and lower viscosity than those of DPPC between 25 and 45 °C (Fig. 3).

**Fig. 2.**
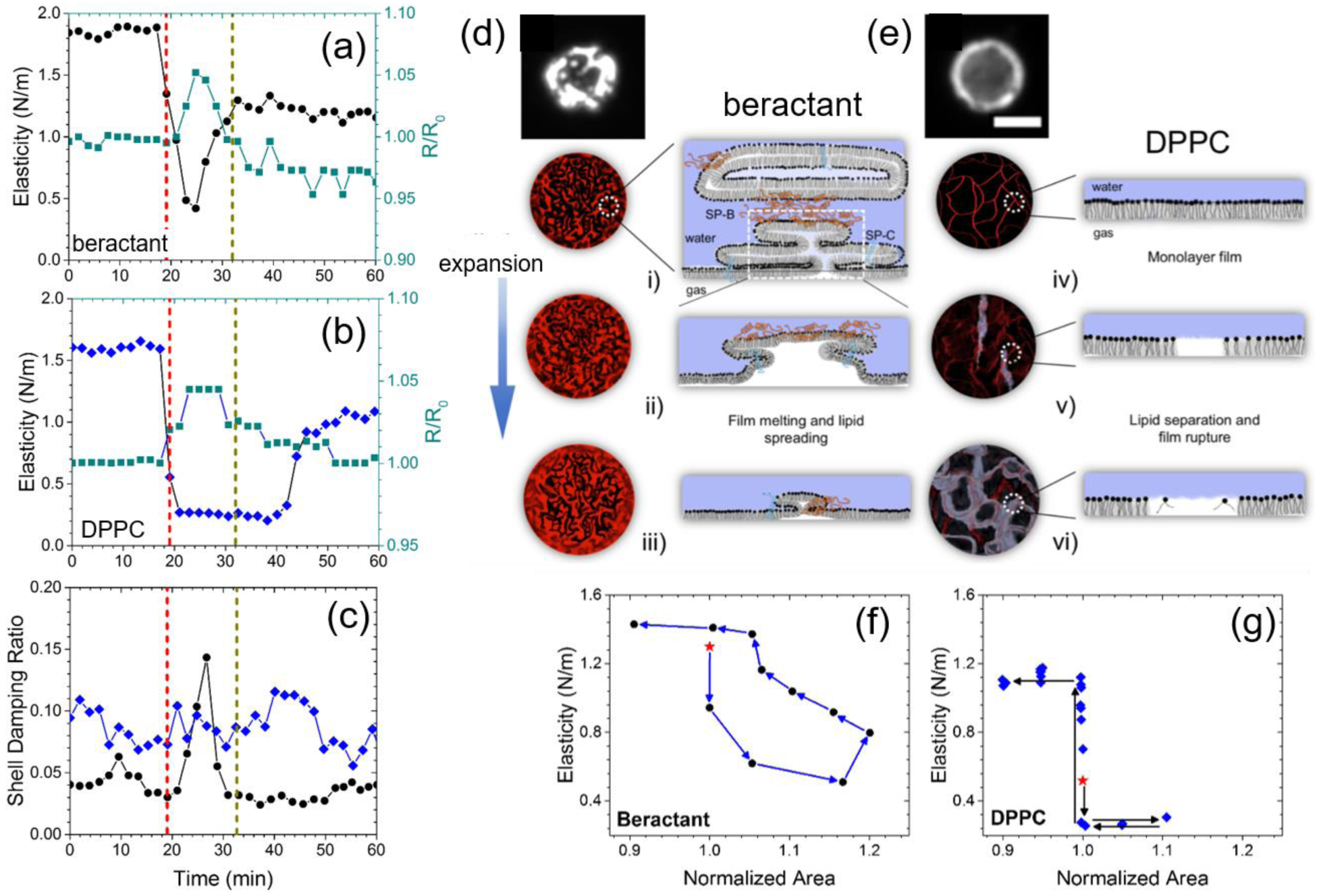
Interfacial melting and spreading stabilizes Beractant films. Representative experiments for individual Beractant (a) and DPPC (b) microbubbles. The red dotted vertical line denotes when heating commenced, and the gray dotted vertical line denotes when body temperature was reached. (c) The shell damping ratio from the same experiments in (a) and (b). Shown are the normalized radius (green squares) and viscoelastic properties for beractant (black circles) and DPPC (blue diamonds). The simultaneous rise in viscosity and drop in elasticity for beractant is a hallmark for interfacial melting and spreading. (d-e) A schematic of the spreading mechanism. Fluorescently labeled beractant (d) and DPPC (e) microbubbles are shown at rest. Beractant microbubbles showed bright, multilayered structures, whereas DPPC microbubbles exhibited a uniform monolayer coating without such multilayers (scale bar 5 μm). (i - iii) The surfactant proteins in lung surfactant help to anchor multilayers onto the surface and facilitate their melting and spreading to incorporate onto the expanding interface, keeping the surface coated. (iv – vi) The DPPC monolayer shell ruptures when the surface expands to expose bare air-water interfaces. Representative plots are shown of the shell elasticity cycle versus normalized area for (f) beractant and (g) DPPC films. The star symbol indicates the first measurement after heating begins, and the arrows indicate the next measurement. Beractant microbubbles showed significant hysteresis, indicating plastic deformation. DPPC microbubbles showed little to no hysteresis, indicating brittle fracture.

**Fig. 3.**
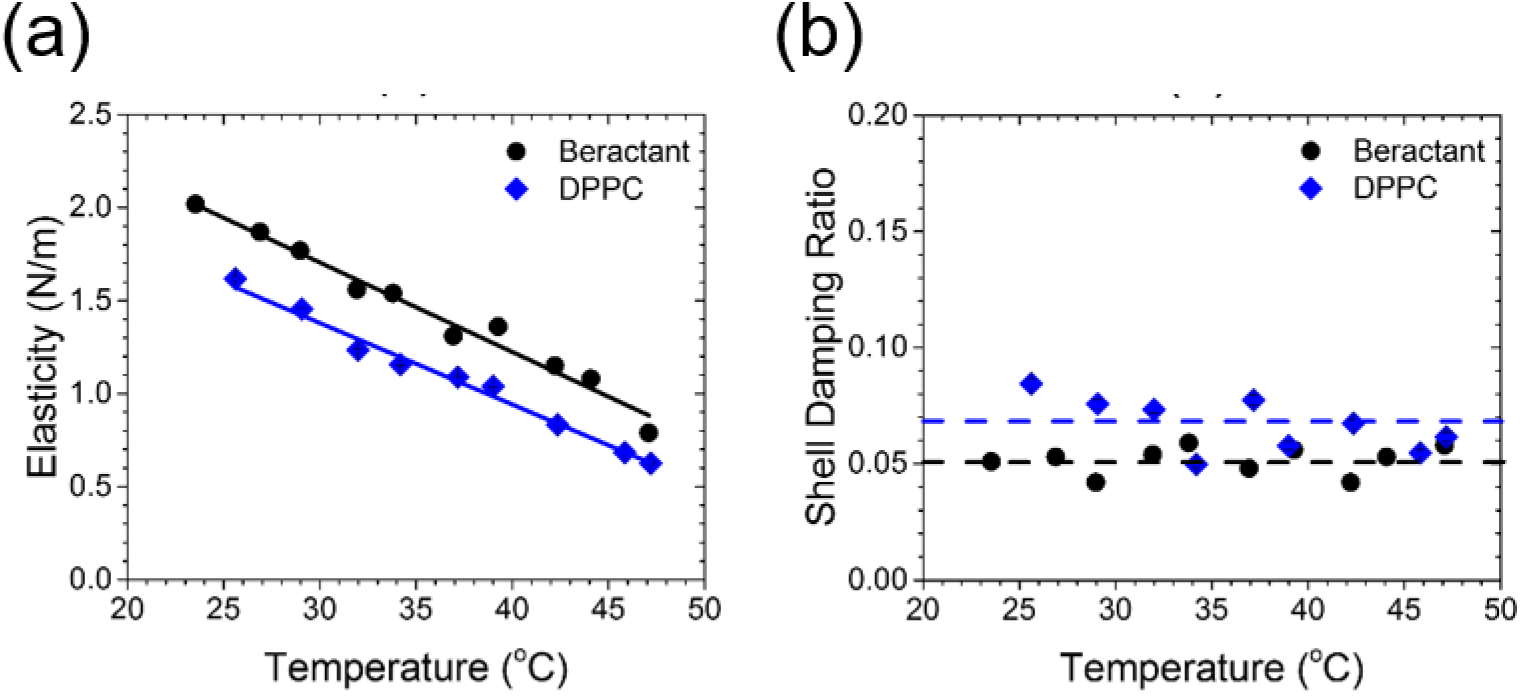
Steady-state viscoelastic properties of the microbubble films. (a) The steady-state dilatational surface elasticity of Beractant and DPPC microbubbles versus temperature (n ≥ 20 for each temperature and shell type). The solid lines are linear fits to the data (Beractant, R^2^ = 0.973; DPPC, R^2^ = 0.986). The elasticity of Beractant was ∼28% greater than that of DPPC. (b) The damping ratio is a measure of shell viscosity and was independent of temperature for DPPC (0.067 ± 0.001) and Beractant (0.052 ± 0.0005). The damping ratio was ∼30% higher for DPPC than Beractant over this temperature range.

We next compared the echogenicity and contrast persistence of vaporized Beractant nanodrops *in vitro* using a medical ultrasound scanner. Nanodrops of decafluorobutane were generated using a microbubble-condensation technique [10]. Beractant was compared to the complete Perflutren lipid formulation, to evaluate against the state of the art. Microbubble size and concentration for Beractant and Perflutren lipid were equalized prior to nanodrop formation by the microbubble-condensation technique (supplementary Table S1 and Fig. S1). The resulting 100-500 nm diameter nanodrops were passed through the focal region of a clinical ultrasound imaging probe (Fig. 4a) and allowed to equilibrate at 37 °C for 5 min. B-mode images showed the onset of ADV as the mechanical index (MI) was increased (Fig. 4b). The threshold MI for the onset of ADV was approximately equal for Beractant and Perflutren lipid (supplementary Fig. S2). However, the echogenicity was higher for Beractant than for Perflutren lipid nanodrops. To quantify echogenicity, the individual pixel video intensities were summed over a region of interest encompassing the tube lumen. The resulting plots show that echogenicity was significantly higher and endured longer for Beractant than Perflutren lipid (Fig. 4c). Indeed, the maximum video intensity was more than twofold higher for Beractant at each MI (Fig. S2b). These results demonstrate that lung surfactant produces phase-change nanodrops with greater echogenicity and stability than the conventional Perflutren lipid formulation, owing to optimal interfacial transformations exhibited by this highly evolved biological material.

**Fig. 4.**
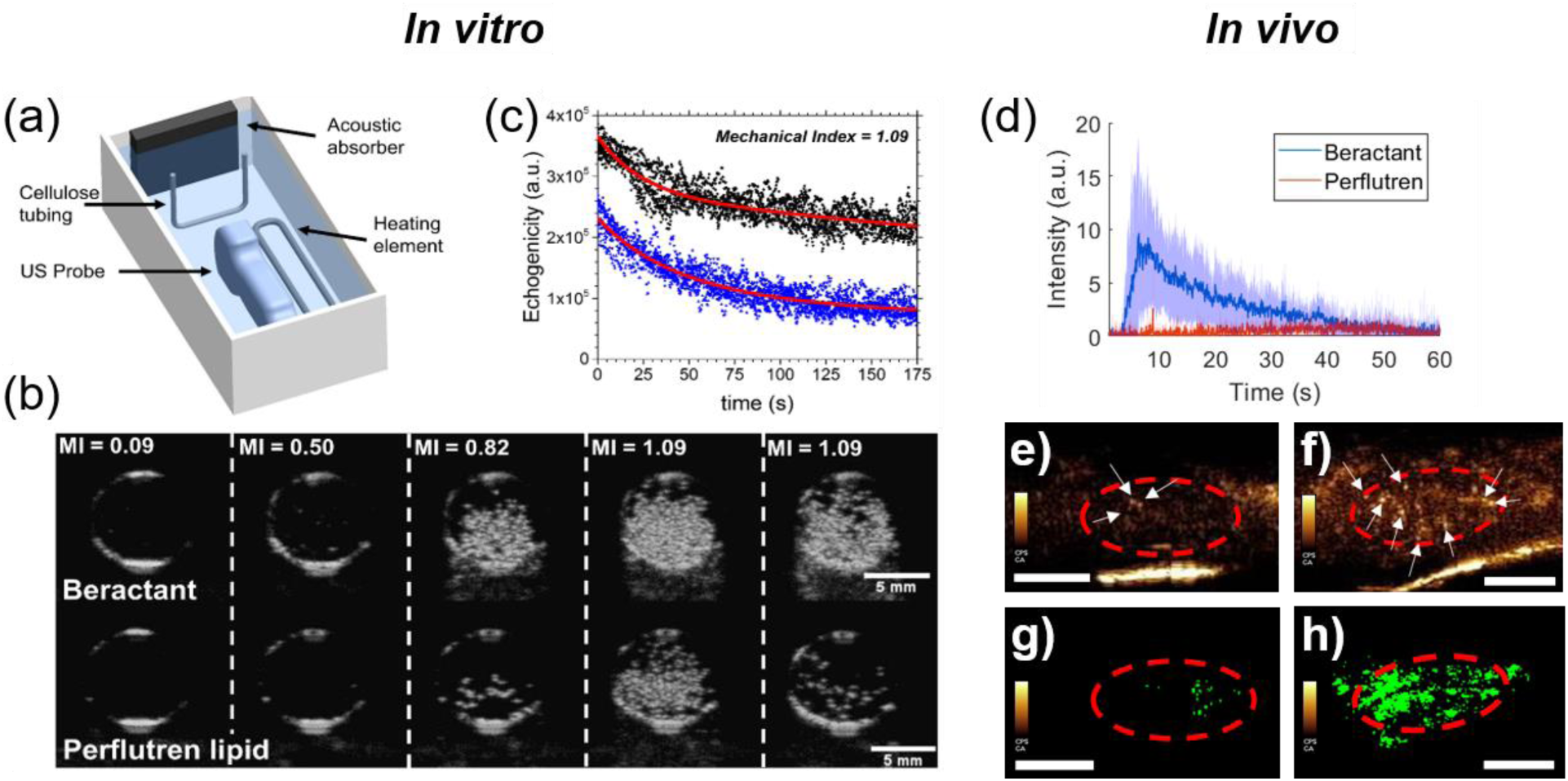
Acoustic droplet vaporization for ultrasound imaging. (a) Experimental diagram showing the cellulose sample chamber and ultrasound probe. (b) B-mode ultrasound images of the cellulose tube cross section taken at 8 MHz at a constant temperature of 37 °C. Representative images of an experiment for Beractant (top panel) and Perflutren lipid (bottom panel) coated nanodrops with a liquid decafluorobutane core. The mechanical index (MI) was increased from 0.09 to 1.09, as shown. (c) The video intensities of six experiments overlaid for Beractant (black, n = 6) and Perflutren lipid (blue, n = 6). The red curves are exponential regressions to guide the eye. In vivo demonstration of contrast generation from Beractant vs. Perflutren lipid nanodrops. (d) Time intensity curves resulting from Beractant and Perflutren lipid nanodrops. Data presented as mean (solid line) ± SD (shaded area) with n = 4 replicates per group. AUCs between the two groups were found significantly different (p<0.0001). The circulation half-life for Beractant was found to be 11.4 s. (e) and (f) Example contrast frames showing microbubbles resulting from Perflutren lipid (e) and Beractant (f) nanodrop vaporization. The region of interest consisting of the kidney is outlined in red and echogenic post-vaporization microbubbles are shown by white arrows. (g) and (h) Example of maximum intensity projection of the kidney vasculature data generated from a single Perflutren lipid (g) and Beractant (h) nanodrop bolus administration using clinically approved contrast doses, hardware and imaging settings. Scale bar = 0.5 cm.

We next demonstrated the utility of lung surfactant nanodrops for *in vivo* vascular imaging. ADV in the mouse kidney was observed for both formulations, but Beractant nanodrops resulted in more contrast as demonstrated by its time-intensity curve (Fig. 4d). Individual contrast pixels were observed in each imaging frame (Fig. 4e,f), and maximum intensity projection images showed perfusion of ADV nanodrops throughout the kidney vasculature for Beractant but not Perflutren lipid (Fig. 4g,h). The poor contrast for Perflutren lipid nanodrops compared to Perflutren microbubbles was likely due to their post-ADV instability, which was a consequence of the relatively poor performance of Perflutren lipids in folding during condensation and re-spreading during ADV. The stochastic activation of nanodrops using ultrasound-imaging pulses at 0.7 MI as done here, combined with the bubble destruction from subsequent imaging pulses, results in ‘activation/deactivation’ signals similarly to the principle described for super-resolution imaging [6]. The Beractant nanodrops allowed us to achieve these images within a single bolus administration in the absence of any motion correction and less than one minute for data collection on a clinical scanner within clinically approved MI, and below the 0.8 MI threshold recently shown to result in detectable bioeffects in the kidney with nanodrop imaging [23]. Thus, the lung surfactant nanodrops show potential for clinical translation of super-resolution imaging.

We next investigated the performance of Beractant nanodrops as CEUS agents for ultrafast ultrasound imaging in a CAM model (Fig. 5). Imaging was conducted using a Verasonics Vantage 256 with an L35-16vX transducer using planewave acquisitions (9-angle coherent compounding, 1-degree step angle, center frequency = 20 MHz). Images were acquired as in-phase quadrature data for a total acquisition length of 1500 frames (30 s). Two different sites in an *ex ovo* chicken embryo, the brain through intact skull and the surface vasculature of the CAM, were examined to visualize resultant microbubbles following ADV, which manifest as individual bright pixels observed in the B-mode images (Fig. 5a,d). A 70-uL bolus of nanodrops was intravascularly injected into a CAM surface microvessel with a glass capillary needle under a dissecting microscope. Using SVD-based filtering [21], ADV nanodrops were readily identified due to their high spatiotemporal coherence as compared to background tissue and blood flow (Fig. 5b,e). Using SVD-based clutter filtering [21], microvessel perfusion images were generated (Fig. 5c,f) with ADV nanodrops superimposed (white arrows). The microvessel images revealed the relative position between tissue microvasculature, and a number of *static* ADV nanodrops, seemingly excluded from visible vasculature. Imaging at 20 MHz, capillaries remain at the threshold of the diffraction/resolution limit, imposing a critical limitation on precise spatial segregation between nanodrops and finer vasculature. Indeed, it is possible that the static nanodrops observed exterior to visible vasculature represented nanodrops trapped within such non-imaged capillaries. However, no evidence of abnormal perfusion, indicative of microvascular occlusion, was observed in nearby, larger vessels, suggesting the possibility that these static nanodrops had extravasated. These activated nanodrops (either extravasated or not) offer opportunities for super-resolution imaging based on localization [7]. When inside the vasculature, nanodrops offer controlled activation and deactivation (by microbubble disruption), which facilitates fast and robust microbubble localization for rapid super-resolution imaging. When outside the vasculature, nanodrops offer the possibility of extravascular super-resolution imaging, which may provide functional assessment of the state of tissue health (e.g., vascular leakiness, molecular imaging with targeted nanodrops etc.).

**Fig. 5.**
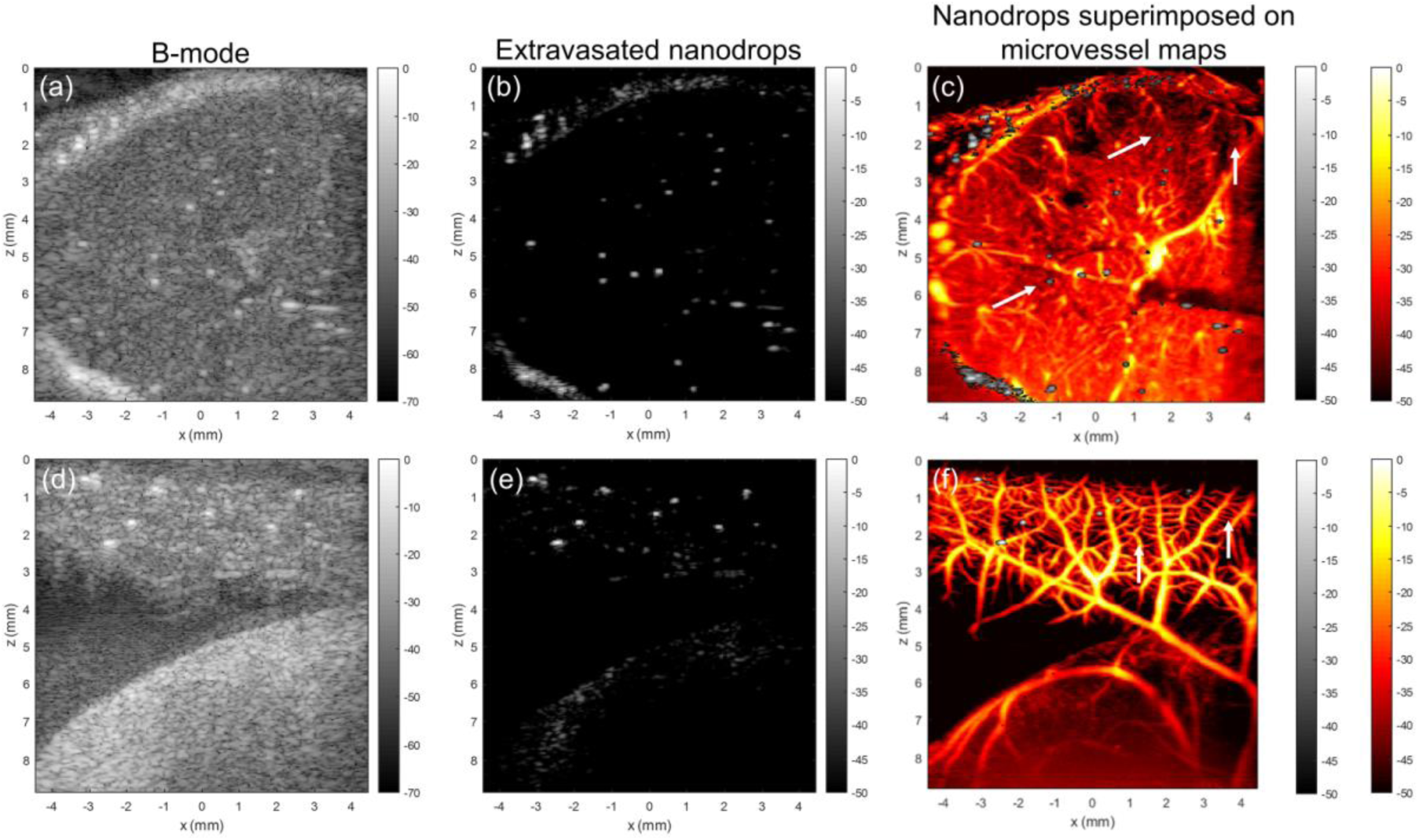
Nanodrop imaging in chicken embryos. a) B-mode ultrasound image of chicken embryo imaged through the intact skull, with ADV nanodrops visible as bright spots. b) Position of vessel-relative position of nanodrops could be isolated in this imaging dataset via singular value decomposition (SVD)-based filtering to extract stationary features. c) SVD clutter filtering was used to generate ultrasound microvessel images. Arrows point to ADV nanodrops. d) B-mode ultrasound image of the chorioallantoic membrane (CAM). e) ADV nanodrop signals isolated using SVD filtering. f) Nanodrop signals overlaid onto ultrasound microvessel perfusion images of CAM vasculature.

Finally, we examined the vaporization and distribution of Beractant nanodrops in 10-week old NOD mice. Vascular leakiness in the pancreas is associated with insulitis in animal models and human Type 1 diabetes (T1D), and can provide an indication of diabetes pathogenesis [24–26]. Detection of diabetes progression is urgently needed to aid in early therapeutic treatment and post-treatment monitoring for T1D prevention. Following Beractant nanodrop infusion and ∼25 min delay to allow for accumulation, nanodrops in both the healthy kidneys and diseased pancreas were stimulated with a small-animal ultrasound scanner, which briefly applied a high intensity acoustic pulse (MI = 1.5) to initiate ADV, then imaged at a lower MI of 0.2. We observed significantly increased contrast intensity in the diseased pancreas but not the kidneys (Fig. 6), demonstrating the enhanced retention of nanodrops in the presence of compromised vasculature.

**Fig. 6.**
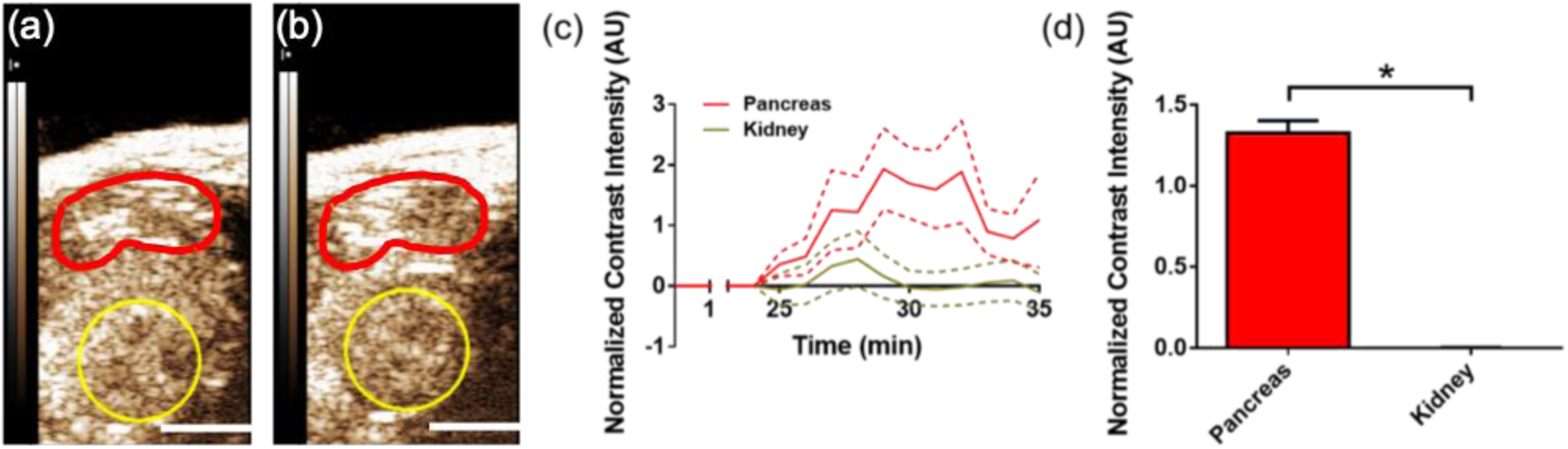
Pancreatic nanodrop accumulation in diabetic mouse model. (a) Ultrasound contrast-mode images in the abdomen of a 10-week old NOD mouse, taken a) pre-ADV, and b) post-ADV of nanodrops. The region of interest indicates the pancreas tail (red) and kidney (yellow; scale bars = 3 mm). (c) Ultrasound contrast intensity post-ADV, normalized to the pre-ADV intensity, as a function of time in the pancreas (green) and kidney (red) [28]. ADV was initiated by a high mechanical index flash of 1.5 at 25 min post nanodrop infusion. Data presented as mean (solid line) ± SD (dashed lines) with n = 6 replicates. (d) mean normalized contrast intensity averaged over time points 30-35 min post-infusion (5-10 min post-ADV). * indicates p < 0.0001 (Student’s t-test).

The lung surfactant nanodrops formed in this study did not spontaneously vaporize at physiological temperature or at very low ultrasound MIs, but rather vaporized in a controllable manner at MIs suitable for diagnostic imaging and therapeutic applications. The implementation of the three different ultrasound systems operating at different transmit frequencies indicate that MI was a suitable parameter to control to initiate ADV, although prior literature has shown inconsistent effects of frequency and pressure on nanodrop ADV [27]. This is an important result illustrating the ability for lung surfactant nanodrops to be rapidly translated to the clinic.

## 4 Conclusions

In summary, we have demonstrated the properties and capabilities of Beractant-encapsulated phase-change nanodrops. Of utility, Beractant is a commercially available and clinically approved lung surfactant replacement therapy. Our results indicate that the Beractant shell provides extreme spreadability through expansion-induced monolayer melting and unfolding of shell-associated folds to coat and stabilize the interface during liquid-to-gas expansion upon vaporization. The expanded Beractant film then rapidly re-solidifies into a stiff shell to stabilize the resultant microbubble. The enhanced stability and echogenicity of post-ADV Beractant microbubbles provides enhanced intravascular imaging, and possible applications of extravascular ultrasound imaging, as suggested in the CAM and T1D mouse models here.

## General

We would like to thank Jordan S. Lum for helpful discussions.

## Author contributions

For design, synthesis and characterization of nanodrops: A.T. and M.B. conceived the research and experiments; T.M. and M.B. supervised the research; A.T. carried out the experiments and analyzed the data. For *in vivo* mouse kidney imaging: A.U. and M.B. conceived the research and experiments; M.B. supervised the research; A.U. and V.P. carried out the experiments and analyzed the data. For *ex ovo* chicken embryo imaging: M.L., P.S. and M.B. conceived the research and experiments; P.S. and M.B. supervised the research; A.U. and M.L. carried out the experiments and analyzed the data. For *in vivo* imaging of the pancreas of the diabetic mouse: D.R. and R.B. conceived the research and experiments; R.B. and M.B. supervised the research; A.U. and D.R. carried out the experiments and analyzed the data. K.H.S., M.B. and A.T. wrote the paper.

## References

[1] S. Wang, J.A. Hossack, A.L. Klibanov, From Anatomy to Functional and Molecular Biomarker Imaging and Therapy: Ultrasound Is Safe, Ultrafast, Portable, and Inexpensive, Invest. Radiol. 55 (2020) 559–572. https://doi.org/10.1097/RLI.0000000000000675.

[2] M.S. Tartis, D.E. Kruse, H. Zheng, H. Zhang, A. Kheirolomoom, J. Marik, K.W. Ferrara, Dynamic microPET imaging of ultrasound contrast agents and lipid delivery, J. Control. Release Off. J. Control. Release Soc. 131 (2008) 160–166. https://doi.org/10.1016/j.jconrel.2008.07.030.

[3] N. Rapoport, K.-H. Nam, R. Gupta, Z. Gao, P. Mohan, A. Payne, N. Todd, X. Liu, T. Kim, J. Shea, C. Scaife, D.L. Parker, E.-K. Jeong, A.M. Kennedy, Ultrasound-Mediated Tumor Imaging and Nanotherapy using Drug Loaded, Block Copolymer Stabilized Perfluorocarbon Nanoemulsions, J. Control. Release Off. J. Control. Release Soc. 153 (2011) 4–15. https://doi.org/10.1016/j.jconrel.2011.01.022.

[4] K. Ferrara, R. Pollard, M. Borden, Ultrasound microbubble contrast agents: fundamentals and application to gene and drug delivery, Annu. Rev. Biomed. Eng. 9 (2007) 415–447. https://doi.org/10.1146/annurev.bioeng.8.061505.095852.

[5] D.G. Ramirez, E. Abenojar, C. Hernandez, D.S. Lorberbaum, L.A. Papazian, S. Passman, V. Pham, A.A. Exner, R.K.P. Benninger, Contrast-enhanced ultrasound with sub-micron sized contrast agents detects insulitis in mouse models of type1 diabetes, Nat. Commun. 11 (2020) 2238. https://doi.org/10.1038/s41467-020-15957-8.

[6] G. Zhang, S. Harput, S. Lin, K. Christensen-Jeffries, C.H. Leow, J. Brown, C. Dunsby, R.J. Eckersley, M.-X. Tang, Acoustic wave sparsely activated localization microscopy (AWSALM): Super-resolution ultrasound imaging using acoustic activation and deactivation of nanodroplets, Appl. Phys. Lett. 113 (2018) 014101. https://doi.org/10.1063/1.5029874.

[7] C. Errico, J. Pierre, S. Pezet, Y. Desailly, Z. Lenkei, O. Couture, M. Tanter, Ultrafast ultrasound localization microscopy for deep super-resolution vascular imaging, Nature. 527 (2015) 499–502. https://doi.org/10.1038/nature16066.

[8] O.D. Kripfgans, J.B. Fowlkes, D.L. Miller, O.P. Eldevik, P.L. Carson, Acoustic droplet vaporization for therapeutic and diagnostic applications, Ultrasound Med. Biol. 26 (2000) 1177–1189. https://doi.org/10.1016/s0301-5629(00)00262-3.

[9] P.A. Mountford, M.A. Borden, On the thermodynamics and kinetics of superheated fluorocarbon phase-change agents, Adv. Colloid Interface Sci. 237 (2016) 15–27. https://doi.org/10.1016/j.cis.2016.08.007.

[10] P.S. Sheeran, S.H. Luois, L.B. Mullin, T.O. Matsunaga, P.A. Dayton, Design of ultrasonically-activatable nanoparticles using low boiling point perfluorocarbons, Biomaterials. 33 (2012) 3262–3269. https://doi.org/10.1016/j.biomaterials.2012.01.021.

[11] P.A. Mountford, W.S. Smith, M.A. Borden, Fluorocarbon nanodrops as acoustic temperature probes, Langmuir ACS J. Surf. Colloids. 31 (2015) 10656–10663. https://doi.org/10.1021/acs.langmuir.5b02308.

[12] Z.Z. Wong, O.D. Kripfgans, A. Qamar, J.B. Fowlkes, J.L. Bull, Bubble evolution in acoustic droplet vaporization at physiological temperature via ultra-high speed imaging, Soft Matter. 7 (2011) 4009–4016. https://doi.org/10.1039/C1SM00007A.

[13] J.J. Kwan, M.A. Borden, Lipid monolayer dilatational mechanics during microbubble gas exchange, Soft Matter. 8 (2012) 4756–4766. https://doi.org/10.1039/C2SM07437K.

[14] S. Sirsi, C. Pae, D.K.T. Oh, H. Blomback, A. Koubaa, B. Papahadjopoulos-Sternberg, M. Borden, Lung surfactant microbubbles, Soft Matter. 5 (2009) 4835–4842. https://doi.org/10.1039/B915065J.

[15] A.N. Thomas, M.A. Borden, Hydrostatic Pressurization of Lung Surfactant Microbubbles: Observation of a Strain-Rate Dependent Elasticity, Langmuir. 33 (2017) 13699–13707. https://doi.org/10.1021/acs.langmuir.7b03307.

[16] J.S. Lum, D.M. Stobbe, M.A. Borden, T.W. Murray, Photoacoustic technique to measure temperature effects on microbubble viscoelastic properties, Appl. Phys. Lett. 112 (2018) 111905. https://doi.org/10.1063/1.5005548.

[17] P.A. Mountford, S.R. Sirsi, M.A. Borden, Condensation phase diagrams for lipid-coated perfluorobutane microbubbles, Langmuir ACS J. Surf. Colloids. 30 (2014) 6209–6218. https://doi.org/10.1021/la501004u.

[18] P.A. Mountford, A.N. Thomas, M.A. Borden, Thermal activation of superheated lipid-coated perfluorocarbon drops, Langmuir ACS J. Surf. Colloids. 31 (2015) 4627–4634. https://doi.org/10.1021/acs.langmuir.5b00399.

[19] D. Chatterjee, K. Sarkar, A Newtonian rheological model for the interface of microbubble contrast agents, Ultrasound Med. Biol. 29 (2003) 1749–1757. https://doi.org/10.1016/s0301-5629(03)01051-2.

[20] G. Montaldo, M. Tanter, J. Bercoff, N. Benech, M. Fink, Coherent plane-wave compounding for very high frame rate ultrasonography and transient elastography, IEEE Trans. Ultrason. Ferroelectr. Freq. Control. 56 (2009) 489–506. https://doi.org/10.1109/TUFFC.2009.1067.

[21] P. Song, A. Manduca, J.D. Trzasko, S. Chen, Ultrasound Small Vessel Imaging With Block-Wise Adaptive Local Clutter Filtering, IEEE Trans. Med. Imaging. 36 (2017) 251–262. https://doi.org/10.1109/TMI.2016.2605819.

[22] C. Huang, M.R. Lowerison, F. Lucien, P. Gong, D. Wang, P. Song, S. Chen, Noninvasive Contrast-Free 3D Evaluation of Tumor Angiogenesis with Ultrasensitive Ultrasound Microvessel Imaging, Sci. Rep. 9 (2019) 1–11. https://doi.org/10.1038/s41598-019-41373-0.

[23] A.G. Nyankima, J.D. Rojas, R. Cianciolo, K.A. Johnson, P.A. Dayton, In Vivo Assessment of the Potential for Renal Bio-Effects from the Vaporization of Perfluorocarbon Phase-Change Contrast Agents, Ultrasound Med. Biol. 44 (2018) 368–376. https://doi.org/10.1016/j.ultrasmedbio.2017.10.016.

[24] Z. Medarova, G. Castillo, G. Dai, E. Bolotin, A. Bogdanov, A. Moore, Noninvasive magnetic resonance imaging of microvascular changes in type 1 diabetes, Diabetes. 56 (2007) 2677–2682. https://doi.org/10.2337/db07-0822.

[25] J.L. Gaglia, M. Harisinghani, I. Aganj, G.R. Wojtkiewicz, S. Hedgire, C. Benoist, D. Mathis, R. Weissleder, Noninvasive mapping of pancreatic inflammation in recent-onset type-1 diabetes patients, Proc. Natl. Acad. Sci. U. S. A. 112 (2015) 2139–2144. https://doi.org/10.1073/pnas.1424993112.

[26] J.R. St Clair, D. Ramirez, S. Passman, R.K.P. Benninger, Contrast-enhanced ultrasound measurement of pancreatic blood flow dynamics predicts type 1 diabetes progression in preclinical models, Nat. Commun. 9 (2018) 1742. https://doi.org/10.1038/s41467-018-03953-y.

[27] M. Aliabouzar, K.N. Kumar, K. Sarkar, Acoustic vaporization threshold of lipid-coated perfluoropentane droplets, J. Acoust. Soc. Am. 143 (2018) 2001. https://doi.org/10.1121/1.5027817.

[28] M.C. Denis, U. Mahmood, C. Benoist, D. Mathis, R. Weissleder, Imaging inflammation of the pancreatic islets in type 1 diabetes, Proc. Natl. Acad. Sci. 101 (2004) 12634–12639. https://doi.org/10.1073/pnas.0404307101.

